# Synthetic auxotrophy remains stable after continuous evolution and in co-culture with mammalian cells

**DOI:** 10.1101/2020.09.27.315804

**Authors:** Aditya M. Kunjapur, Michael G. Napolitano, Eriona Hysolli, Karen Noguera, Evan M. Appleton, Max G. Schubert, Michaela A. Jones, Siddharth Iyer, Daniel J. Mandell, George M. Church

**Author notes:** Corresponding authors: AMK or GMC. These authors contributed equally to this work (Aditya M. Kunjapur, Michael G. Napolitano, Eriona Hysolli).

## Abstract

Understanding the evolutionary stability and possible context-dependence of biological containment techniques is critical as engineered microbes are increasingly under consideration for applications beyond biomanufacturing. While batch cultures of synthetic auxotrophic *Escherichia coli* previously exhibited undetectable escape throughout 14 days of monitoring, the long-term effectiveness of synthetic auxotrophy is unknown. Here, we report automated continuous evolution of a synthetic auxotroph using custom chemostats that supply a decreasing concentration of essential biphenylalanine (BipA). After 100 days of evolution in three separate trials, populations exhibit no observable escape and are capable of normal growth rates at 10-fold lower BipA concentration than the ancestral synthetic auxotroph. Allelic reconstruction of three proteins implicated in small molecule transport reveals their contribution to increased fitness at low BipA concentrations. Mutations do not appear in orthogonal translation machinery nor in synthetic auxotrophic markers. Based on its evolutionary stability, we introduce the progenitor synthetic auxotroph directly to mammalian cell culture. We observe containment of bacteria without detrimental effects on HEK293T cells. Overall, our findings reveal that synthetic auxotrophy is effective on timescales and in contexts that enable diverse applications.

**One Sentence Summary:** To ascertain whether life inevitably finds a way, we continuously evolve an *Escherichia coli* strain that was not able to escape from engineered biocontainment before, and we find that it does not escape even after 100 days of evolution, nor does it escape when added to mammalian cell culture.

## Introduction

New safeguards are needed for the deliberate release of engineered microbes into the environment, which has promise for applications in agriculture, environmental remediation, and medicine^1^. Genetically encoded biocontainment strategies enable attenuation of engineered live bacteria for diverse biomedical applications^2–4^, including as potential vaccines^5–10^, diagnostics^11^, and therapeutics^12–15^. Auxotrophy, which is the inability of an organism to synthesize a compound needed for its growth, is an existing strategy for containment. However, foundational studies of auxotrophic pathogens demonstrated proliferation in relevant biological fluids^16^ and reversion to prototrophy upon serial passaging^17,18^. Modern genome engineering strategies can prevent auxotrophic reversion, and auxotrophy has been a key component of microbial therapies that have reached advanced clinical trials. However, the ability for auxotrophs to access required metabolites within many host microenvironments, and after leaving the host, remains unaddressed. Auxotrophy may not be effective in scenarios where engineered living bacteria encounter metabolites from dead host cells^19^ or invade host cells^20^. Indeed, growth of double auxotrophs is supported *in vivo* by neoplastic tissue^13^. Auxotrophy may also be insufficient for tight control of cell proliferation in environments rich with microbial sources of cross-feeding^21^, such as gut, oral, skin, and vaginal microbiomes. Given that most naturally occurring microorganisms are auxotrophs^22^, it is also unlikely that auxotrophy will limit the spread of an engineered microbe once it leaves the body and enters the environment.

Synthetic auxotrophy may overcome these hurdles by requiring provision of a synthetic molecule for survival of the engineered bacteria. This strategy was first implemented successfully in *E. coli* by engineering an essential protein to depend on incorporation of a non-standard amino acid (nsAA)^23,24^. We previously engineered *E. coli* strains for dependence on the nsAA biphenylalanine (BipA) by computer-aided redesign of essential enzymes in conjunction with expression of orthogonal translation machinery for BipA incorporation^23^. Among several synthetic auxotrophs originally constructed, one strain harbored three redesigned, nsAA-dependent genes: Adenylate kinase (*adk.d6*), tyrosyl-tRNA synthetase (*tyrS.d8*), and BipA-dependent aminoacyl-tRNA synthetase, for aminoacylation of BipA (*BipARS.d6*). This BipA-dependent strain, dubbed “DEP”, exhibited undetectable escape throughout 14 days of monitoring at an assay detection limit of 2.2×10^−12^ escapees per colony forming unit (CFU)^23^. Though this strain demonstrates effective biocontainment in 1 L batch experiments, its precise escape frequency and long-term stability remained unexplored.

Here, we perform the first study of evolutionary stability of a synthetic auxotroph, with the aid of automated continuous evolution. Continuous evolution better emulates scenarios where biocontainment may be needed by fostering greater genetic variability within a population. We posited that decreasing BipA concentrations would add selective pressure for adaptation or for escape, either of which would be enlightening. Adaptive laboratory evolution of DEP may improve its fitness in relevant growth contexts, as previously demonstrated for its non-auxotrophic but recoded ancestor, C321.ΔA^25^. We report that DEP maintains its inability to grow in the absence of synthetic nutrient, even after three parallel 100-day chemostat trials. Additionally, we find evidence of adaptation, with evolved DEP isolates requiring 10-fold lower BipA concentration to achieve optimal growth than ancestral DEP (0.5 μM rather than 5 μM). We resequence evolved populations and perform allelic reconstruction in ancestral DEP using multiplex automatable genome engineering (MAGE), identifying alleles that partially restore the adaptive phenotype. Finally, we advance this technology towards host-microbe co-culture applications, demonstrating direct mixed culture of DEP and mammalian cells without need for physical barriers nor complex fluidics.

## Results

### Continuous evolution

To perform continuous evolution of *E. coli*, we constructed custom chemostats for parallelized and automated culturing (**Fig. 1A**). Our design and construction was based on the eVOLVER system^26^, an open-source, do-it-yourself automated culturing platform (**Figs. S1-S4**). By decreasing BipA concentration over time in our chemostats, we provide an initial mild selection for escape and steadily increase its stringency. This design is analogous to a “morbidostat”, where a lethal drug is introduced dynamically at sub-lethal concentrations to study microbial drug resistance^27^, but with synthetic auxotrophy providing selective pressure. Our working algorithm for automated adjustment of BipA concentration as a function of turbidity is shown in **Fig. 1B**, and a representative image of our hardware is shown in **Fig. 1C.**

**Figure 1.**
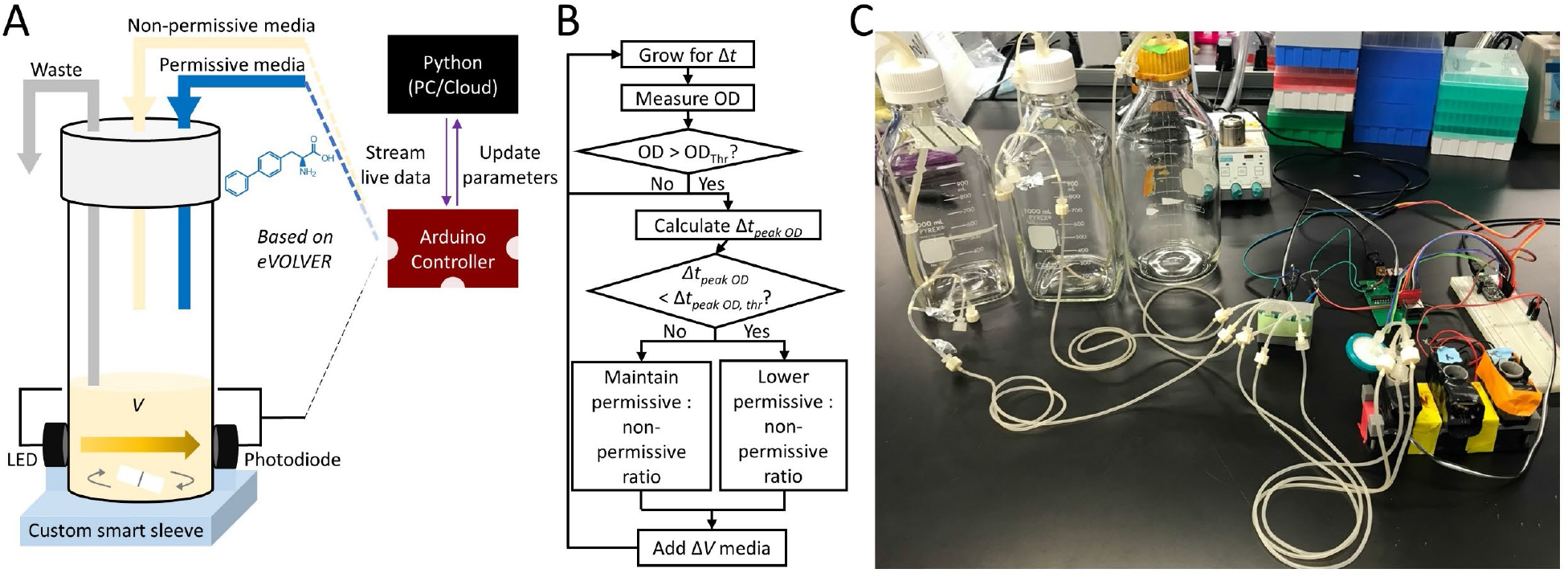
Design and operation of custom chemostats for automated continuous evolution. (A) Illustration of a smart sleeve connected to separate non-permissive media and biphenylalanine (BipA, structure shown in blue) feed lines for automated adjustment of BipA concentration based on growth rate. Pumps and optics are integrated with Arduino controller hardware and Python software based on the eVOLVER do-it-yourself automated culturing framework. (B) Working algorithm for maintenance of cultures in continuous evolution mode. Criteria for lowering the BipA concentration is based on the difference in time elapsing between OD peaks (Δ*t_peak OD_*). Smaller time elapsed between OD peaks is indicative of higher growth rates, triggering decrease of BipA concentration when below a threshold value. (C) Representative configuration of hardware for parallelized evolution in triplicate, with three empty sleeves shown.

We inoculated one chemostat with DEP for an 11-day incubation at constant BipA concentration (100μM) (**Fig. 2A**). After observing no colony formation on non-permissive media, we then inoculated three chemostats in parallel where BipA supply decreased automatically over the following 90 days from 100 μM to nearly 10 nM (**Fig. 2B**). To determine whether the decrease in BipA supply was due to escape from dependence on BipA, we periodically performed escape assays. We continued to observe no escape, including when we seeded liter-scale cultures and plated the associated outgrowth on non-permissive media. Evolved isolates were obtained after this procedure, and their growth was characterized across BipA concentrations (**Fig. 2C**). At 0.5-1 μM BipA, we observed growth of all evolved isolates, and no growth of the ancestral DEP strain.

**Figure 2.**
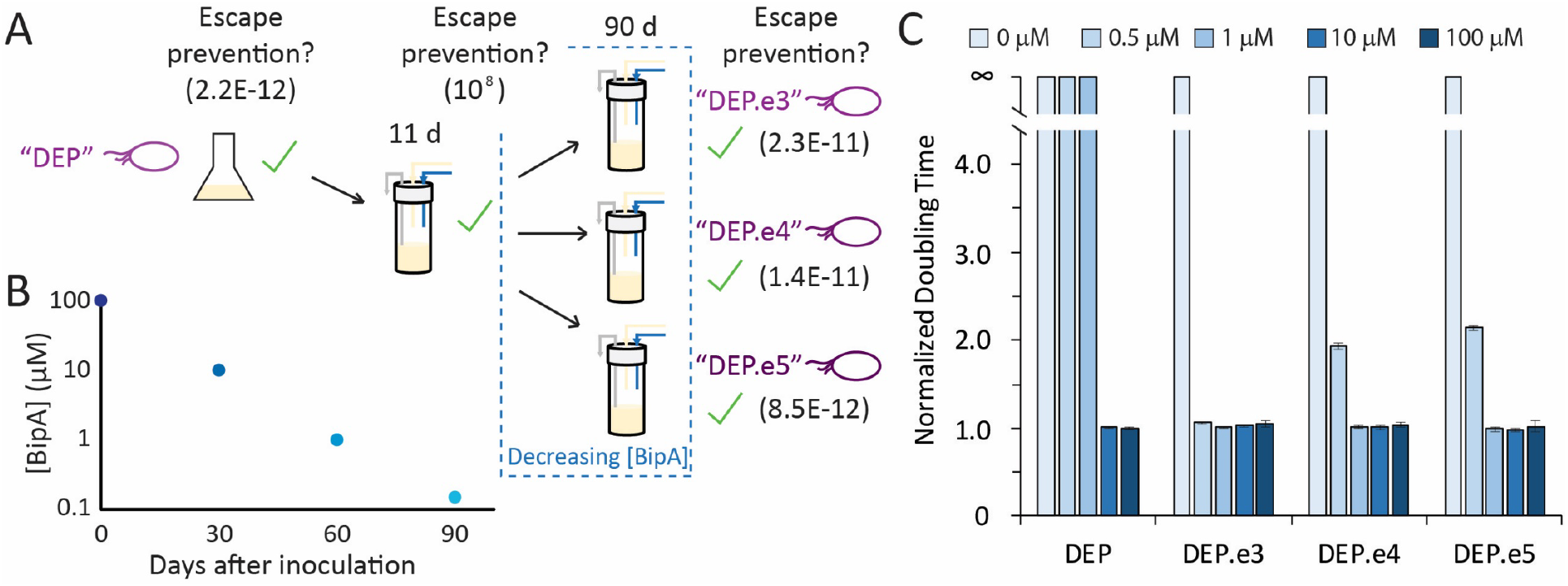
Continuous evolution of synthetic auxotrophs leads to adaptation to lower BipA concentration rather than escape. (A) Timeline for continuous evolution, with detection limits for escape frequency assays shown in parentheses. (B) Concentration of BipA provided to cells during evolution over a 90-day period. (C) Doubling times of progenitor and evolved synthetic auxotrophs as a function of BipA concentration, normalized to the doubling time of DEP at 100 μM BipA.

### Genome resequencing and allelic reconstruction

To identify the causal alleles contributing to decreased BipA requirement of all three evolved isolates, we performed whole-genome sequencing and mutational analysis. We expected that mutations in auxotrophic markers or orthogonal translation machinery associated with aminoacylation of BipA would be observed. However, no variants were detected in the plasmid-expressed orthogonal translation machinery (aminoacyl-tRNA synthetase and tRNA) reference sequence. Instead, in all three evolved isolates, variants were observed in three non-essential genes, all of which are implicated in molecular transport: *acrB. emrD*, and *trkH* (**Fig. 3A**). AcrB and EmrD are biochemically and structurally well-characterized multi-drug efflux proteins^28^, and TrkH is a potassium ion transporter^29^. These exact mutations have no precedent in the literature to our knowledge. Because they are missense mutations or in-frame deletions, it is unclear whether they cause loss of function or altered function (**Table S1**). Because permissive media contains four artificial targets of efflux (BipA, L-arabinose, chloramphenicol, and SDS), mutations that confer a selective advantage during continuous evolution could disable BipA/L-arabinose efflux, improve chloramphenicol/SDS efflux, or affect transport of these or other species more indirectly. Given the strong selective pressure enforced by decreasing BipA concentration, we hypothesize that mutations observed are more likely to affect BipA transport. We also observed mutations in all evolved populations to the 23S ribosomal RNA (rRNA) gene *rrlA* (**Table S2**). 23S rRNA mutations have been found to enhance tolerance for D-amino acids^30^ and β-amino acids^31^. However, 23S rRNA mutations could also be related to increased tolerance of chloramphenicol^32^.

**Figure 3.**
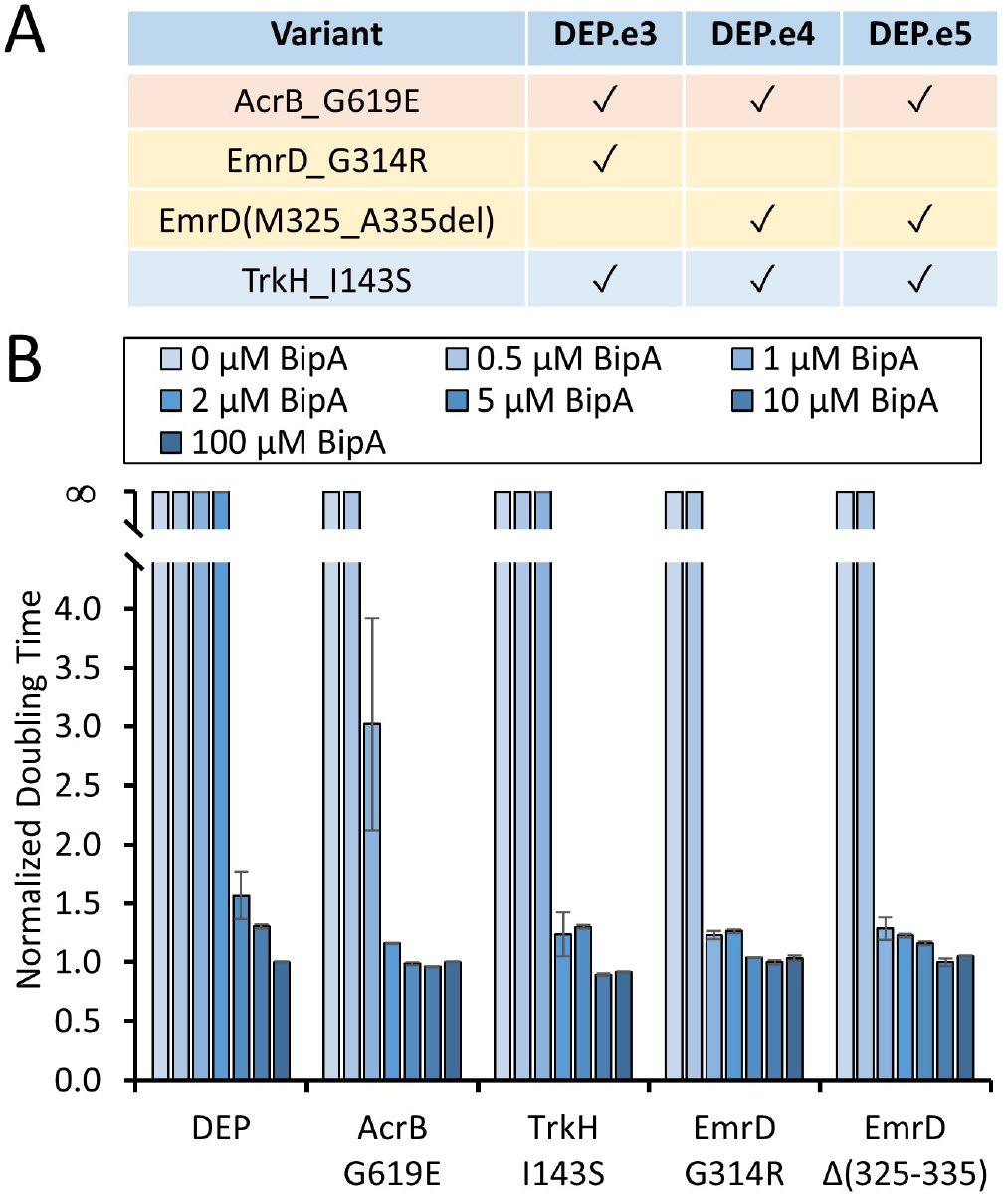
Whole genome sequencing and reconstruction of alleles shared across populations of evolved synthetic auxotrophs. (A) List of alleles identified through next-generation sequencing. (B) Effect of reconstructed allele in DEP progenitor on doubling time as a function of BipA concentration.

To learn how identified transporter alleles may contribute to increased growth rates at low BipA concentration, we performed allelic reconstruction in the progenitor DEP strain using multiplexed automatable genome engineering (MAGE)^33^. Among four single mutants that we generated in DEP, which included both observed *emrD* mutant alleles, we observed growth of all mutants rather than DEP at 2 μM BipA (**Fig. 3B**). Furthermore, only *emrD* mutants exhibited near-normal growth at 1 μM BipA. To investigate possible differential sensitivity of strains that contain reconstructed alleles to other media components of interest (SDS, L-arabinose, Tris buffer, and Chloramphenicol), we varied the concentration of these components and measured doubling times (**Fig. S7**). We observed no significant deviation in doubling time from DEP in any of these cases. These results collectively suggest that observed transporter alleles are linked to BipA utilization.

### Bacterial-mammalian co-culturing

The unobservable escape of DEP even after 100 days of evolution encouraged us to explore the possibility of an improved *in vitro* model for host-microbe interactions. *In vitro* models allow direct visualization and measurement of cells and effectors during processes such as pathogenesis^34^. They are more relevant than animal studies for several human cell-specific interactions due to biological differences across animal types^35,36^. A non-pathogenic *E. coli* strain engineered to express heterologous proteins could be particularly useful for studying or identifying virulence factors and disease progression. However, an obstacle associated with co-culture of microbial and mammalian cells is microbial takeover of the population. Approaches used to address this are bacteriostatic antibiotics^37^, semi-permeable Transwell membranes^38–40^, microcarrier beads^41^, microfluidic cell-trapping^42^, peristaltic microfluidic flow^43,44^, and microfluidic perfusion^45^. However, the use of a well-characterized synthetic auxotroph capable of limited persistence could offer a superior alternative for spatiotemporal control of microbial growth, especially for studying longer duration phenomena such as chronic infection or wound healing.

We investigated mammalian cell culture health, growth, and morphology after simple transient exposure to a hypermutator variant of DEP that we engineered by inactivating *mutS* during allelic reconstruction (DEP*). The use of DEP* rather than DEP is yet another form of a stress test to increase opportunity for escape under co-culture conditions. We directly co-cultured adherent human cell line HEK293T with either no bacteria, non-auxotrophic *E. coli* DH5α, or DEP* overnight (24 hours). HEK293T cells were cultured in selection media that allow only growth of desired but not contaminant strains while selecting for bacterial plasmid maintenance. After co-culture, we washed cells, and replenished cells with media varying in inclusion of BipA and/or an antibiotic cocktail (Penicillin/Streptomycin/Amphotericin B). We continued incubation and imaged cells at Days 2, 4, and 7 after initial co-incubation. HEK293T cells contain a copy of mCherry integrated into the AAVS1 locus and they appear red. DH5α and DEP* were transformed with Clover green fluorescent protein prior to co-culture and appear green.

Compared to the control culture where bacteria were not added (**Fig. 4A**), HEK293T cells co-cultured with DH5α display visible bacterial lawns with no attached human cells in the absence of the antibiotic cocktail at all days of observation (**Fig. 4B**). In the presence of antibiotic, co-cultures containing DH5α sharply transition from bacterial overgrowth to apparent bacterial elimination (**Fig. 4C**). In contrast, cells co-cultured with DEP* in the absence of BipA exhibited similar morphology to the control at all days of observation and no detectable bacteria by fluorescence microscopy on Day 7, without the need for antibiotics to achieve bacterial clearance (**Fig. 4D**). Thus, DEP* addition was not detrimental to HEK293T cells in the absence of BipA and DEP* remains biocontained and cannot survive because of cross-feeding. Clearance of bacterial cells from human cells appears to occur faster for DEP* when not provided BipA (**Fig. 4D**) than for DH5α when provided the antibiotic cocktail (**Fig 4C**).

**Figure 4.**
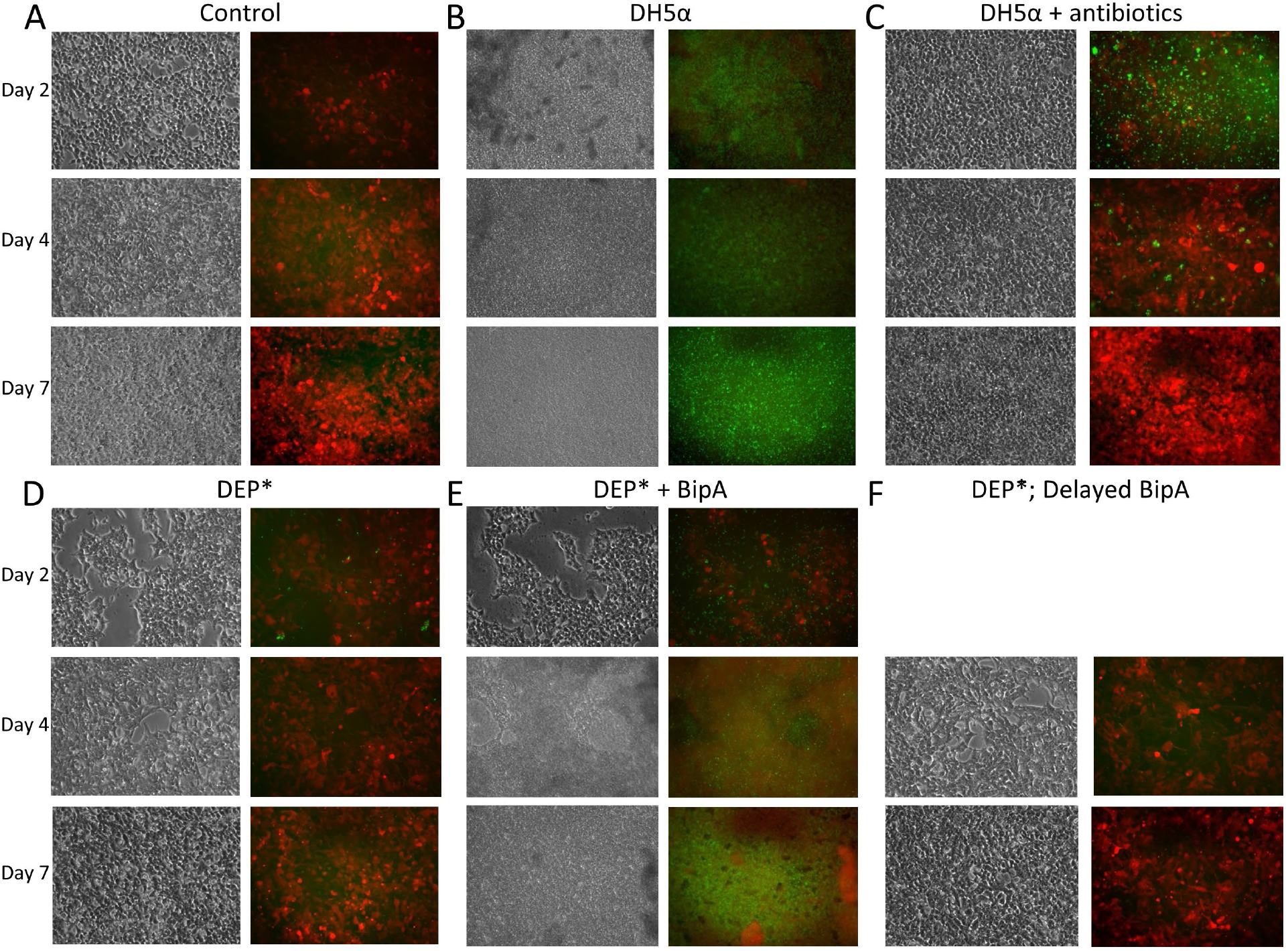
Imaging of bacterial-mammalian co-cultures. Bacteria were added to HEK293T cell cultures and co-incubated for 24 hours before washing and replenishing media. HEK293T cells express mCherry, whereas bacterial cells express Clover green protein marker. Images were taken at Days 2, 4, and 7 after co-incubation. (A) Untreated HEK293T cells. (B) HEK293T with commercial *E. coli* DH5α in the absence of antibiotic cocktail. (C) HEK293T with DH5α in presence of antibiotic cocktail. (D) HEK293T and DEP*\** (mismatch repair inactivated to create hypermutator phenotype) in the absence of BipA. (E) HEK293T cells and DEP*\** in the presence of BipA. (F) HEK293T and DEP* in the absence of BipA until Day 2 (identical at this point to condition in Panel D), and then 100 μM of BipA added to this condition daily until Day 7.

To learn how the synthetic auxotroph behaves when supplied its essential nutrient in these co-culture settings, we tested DEP* co-cultures with continual resupply of 100 μM BipA. Here, DEP* proliferates and in turn decreases proliferation and viability of HEK293T cells (**Fig. 4E**). A bacterial lawn begins to form on Day 2 and at later times human cell debris is overtaken by DEP*. This demonstrates that DEP* is fully capable of taking over the co-culture if supplied with BipA. Replicates for these experiments can be found in **Figs. S8-10**.

### Persistence

Given that DEP* grows in co-cultures when BipA is provided, we sought to understand whether it could be rescued by re-addition of BipA after multiple days of withholding. The possible timescale of re-emergence influences applications where the duration of bacterial activity would need to be prolonged and/or repeated via limited BipA introduction while remaining contained. We find that co-culturing DEP* with HEK293T cells for 2 days in absence of BipA followed by addition of BipA at Day 2 does not rescue the DEP* growth (**Fig. 4F and S11**). Human cells still grow and look morphologically similar to untreated cells and bacteria are not visible. To look at analogous questions for non-auxotrophic *E. coli*, we removed antibiotics after 2 days of co-culturing and do not observe bacterial rescue (**Fig. S11**). We also investigated whether bacterial clearance could be delayed by addition of antibiotic after some growth of DH5α. DH5α cells grown in absence of the antibiotic cocktail for 2 days before addition of the cocktail and maintenance to Day 7 result in bacterial lawns (**Fig. S11A and D**). This demonstrates that antibiotic cocktails ordinarily used in mammalian cell culture maintenance can become ineffective beyond a certain amount of non-auxotrophic bacterial growth, whereas synthetic auxotrophy is subject to fewer and different constraints.

To further investigate the persistence of progenitor DEP and its evolved descendants, we performed BipA re-addition studies in LB monoculture. Within 7 hours of BipA removal, DEP cell populations that are harvested from mid-exponential or stationary phases can be “reactivated” upon delayed BipA addition with unperturbed growth kinetics after a highly tunable lag phase (**Fig. S12**). Growth curve and colony count experiments for cells deprived of BipA for longer timescales demonstrate that growth can be recovered upon BipA reintroduction after as long as 48 hours under these monoculture conditions (**Figs. S13-14**).

We have shown that synthetic auxotrophy can exhibit long-term stability and function in unique contexts, enabling reliable control of microbial proliferation. Recent work has also shown that the escape rate and fitness of multiple synthetic auxotrophs can be improved by increasing the specificity of nsAA incorporation machinery^46^. Collectively, these engineering and characterization efforts advance synthetic auxotrophy as a powerful safeguard for basic and applied research when using engineered microbes.

## Acknowledgements

We thank Caleb Bashor for assistance setting up the continuous evolution platform, and Sabyasachi Sen and Neil Butler for manuscript comments. We thank Timothy Wannier for sharing the Clover plasmid, and Ellen Schrock for helping generate the mCherry stable HEK293T cell line. This work was primarily funded by grants to GMC from the U.S. Dept. of Energy (FG02-02ER63445) and the FunGCAT program from the Office of the Director of National Intelligence (ODNI), Intelligence Advanced Research Projects Activity (IARPA), via the Army Research Office (ARO) under Federal Award No. W911NF-17-2-0089. We also acknowledge support from the National Institute of General Medical Sciences of the National Institutes of Health under a Chemistry-Biology Interface Training Grant that supported MAJ (Award Number T32GM133395), and a grant to AMK from the National Science Foundation (MCB2027092).

The authors declare the following competing financial interest(s): GMC has related financial interests in 64-x, EnEvolv, and GRO Biosciences. For a complete list of GMC’s financial interests, please visit arep.med.harvard.edu/gmc/tech.html.

