## Supplementary Information for "Synthetic auxotrophy remains stable after continuous evolution and in co-culture with mammalian cells"

**Materials and Methods**

**SI Figures 1-14**

**SI Tables 1-4**

**SI References**

### Materials and Methods

#### *Culture conditions*

Cultures for general culturing, growth rate assays, biocontainment escape assays, MAGE, and fluorescent protein assays were prepared in LB-Lennox medium (LB$^L$: 10 g/L bacto tryptone, 5 g/L sodium chloride, 5 g/L yeast extract) supplemented with 15 µg/mL chloramphenicol, 0.2% (wt/v) L-arabinose, 20 mM Tris-HCl buffer, 0.005% SDS, and variable concentration of *L*-4,4-Biphenylalanine (BipA). Unless otherwise indicated, all cultures were grown in biological triplicate in 96-well deep plates in 300 µL culture volumes at 34 °C and 400 rpm. The above media is permissive for growth of the synthetic auxotroph. Non-permissive media is identically formulated as permissive media except for BipA, which is not included.

#### *Construction of custom chemostats*

Construction of appropriate fluidics and chambers followed the eVOLVER framework[1] (**Figs. S1 and S2)**. The following components were included: (1) Fluidics and chambers (reactor vial, inlet and outlet lines, filters, pumps, stirrers, and inlet and outlet reservoirs); (2) Light source and detector (LED and photodiode); (3) Controller hardware (circuit and microprocessors); (4) Controller software (Arduino for controlling tasks, Raspberry Pi for computing tasks, Python code for programming tasks) (full Build of Materials included in **Table S3**). Briefly, our apparatus consisted of a custom "smart sleeve" (**Fig. S3**), with the following modifications: Each vial was constructed without temperature control and was supplied by two media pumps (one for permissive media, another for non-permissive media) and connected to one waste pump. All pumps were RP-Q1 from Takasago fluidics, each driven off a standard N power MOSFET with an Arduino controlling the gate. Like the eVOLVER system, we installed a stirring fan underneath each sleeve that consisted of magnets attached to a computer fan. By including a small stir bar within each reactor vial, we enabled efficient mixing of 1 mL working volumes. To enable automated measurement of turbidity (optical density, or OD), we used a 605 nm LED (LO Q976-PS-25) and an OPT101P-J photodiode detector. We mounted the LED and detector on custom PCBs mounted to the vial sleeve to enable easier construction and better control of ambient light leakage into the light path (**Fig. S4**). To monitor turbidity within each vial and to control pump arrays in response, we constructed printed circuit board designs in Gerber format as is standard for circuit fabrication. We attached an Arduino Mega microcontroller with an Analog-Digital Converter and directed it using a PyMata script.[2]

#### *Operation of custom chemostats*

Chemostats were operated by automated maintenance of culture OD within a specified parameter range within exponential growth phase (20-80% of dynamic range) depending on linearity of photodiode measurements. Constant fixed dilutions of permissive media were used to decrease OD until desired equilibrium of cell growth and dilution rates. This resulted in a sawtooth curve[3], where time between peaks is recorded as a proxy for growth rate. Our program gradually decreased the ratio of permissive to non-permissive media as step functions, with a specified number of dilution cycles allowed to elapse before the next decrease to provide time for acclimation. Time between OD peaks lengthened as strain fitness decreased. Once a threshold difference between ancestral peak-to-peak time and current peak-to-peak time was passed, the ratio of permissive to non-permissive media remained fixed. This allowed cells to evolve until peak-to-peak time returns to ancestral values, which initiated the next phase of decrease in BipA concentration. To assess the quality of our continuous evolution process, we paused chemostat trials on a weekly basis for strain storage, strain evaluation, chemostat cleaning, and investigation of contamination.

*Measurement of doubling times*

Growth assays were performed by plate reader with blanking as previously described[4]. Overnight cultures were supplemented with different BipA concentrations depending on the strain. The DEP progenitor strain was grown in permissive media containing 100 μM BipA, and evolved DEP strains DEP.e3, DEP.e4, and DEP.e5 were grown in permissive media containing 1 μM BipA. Saturated overnight cultures were washed twice in LB and resuspended in LB. Resuspended cultures were diluted 100-fold into three 150 μL volumes of permissive media. BipA concentrations used in this assay were: 0 μM, 0.001 μM, 0.01 μM, 0.1 μM, 0.5 μM, 1 μM, 10 μM, and 100 μM. Cultures were incubated in a flat-bottom 96-well plate (34 °C, 300 r.p.m.). Kinetic growth ($OD_{600}$) was monitored in a Biotek Eon H1 microplate spectrophotometer reader at 5-min intervals for 48 h. The doubling times across technical replicates were calculated as previously indicated.

*Escape frequency assays*

Escape assays were performed as previously described with minor adjustments to decrease the lower detection limit for final evolved populations[5,6]. Strains were grown in permissive media and harvested in late exponential phase. Cells were washed twice with LB and resuspended in LB. Viable CFU were calculated from the mean and standard error of the mean (SEM) of three technical replicates of tenfold serial dilutions on permissive media. Twelve technical replicates were plated on noble agar combined with non-permissive media in 500 $cm^2$ BioAssay Dishes (Thermo Scientific 240835) and monitored daily for 4 days. If synthetic auxotrophs exhibited escape frequencies above the detection limit (lawns) on non-permissive media, escape frequencies were calculated from additional platings at lower density. The SEM across technical replicates of the cumulative escape frequency was calculated as previously indicated.

*Genome resequencing and analysis*

Genomic DNA was obtained from evolved populations and ancestral clone using the Wizard Genomic DNA purification kit (Promega). Sequencing libraries were prepared as described in Baym et al.[7] Sequencing was performed using a NextSeq instrument, producing 75 bp, paired-end reads. Resulting data was aligned to the *E. coli* C321.delA non-auxotrophic but recoded reference sequence (Genbank CP006698.1) and the sequence of the plasmid encoding nsAA incorporation machinery. The Millstone software suite was used to identify variants, provide measures of sequencing confidence, and predict their likelihood of altering gene function[8]. Genomic variants of low confidence, low sequence coverage, or presence in the ancestral strain were discarded, prioritizing variants observed in three non-essential genes that encode membrane proteins: *acrB*. *emrD*, and *trkH*.

*Allelic reconstruction*

Multiplex Automatable Genomic Engineering (MAGE)[9] was used to inactivate the endogenous *mutS* gene in the DEP strain. Overnight cultures were diluted 100-fold into 3 mL $LB^L$ containing chloramphenicol, BipA, L-arabinose, and Tris HCl buffer and grown at 34 °C until mid-log. The genome-integrated lambda Red cassette in this C321.ΔA-derived strain was induced in a shaking water bath (42 °C, 300 rpm, 15 minutes), followed by cooling the culture tube on ice for at least two minutes. The cells were made electrocompetent at 4°C by pelleting 1 mL of culture (8,000 rcf,

30 seconds) and washing thrice with 1 mL ice-cold 10% glycerol. Electrocompetent pellets were resuspended in 50µL of $dH_2O$ containing the desired DNA, for MAGE oligonucleotides, 5 µM of each oligonucleotide was used. Allele-specific colony PCR was used to identify desired colonies resulting from MAGE as previously described[10]. Oligonucleotides used for MAGE and for allele-specific colony PCR are included in **Table S4**.

*Investigation of media conditions on reconstructed alleles*

This assay was performed using a similar protocol as described in the *Measurement of doubling times* section. The cultures for DEP and its single mutants were grown overnight in 100 µM BipA. Then cultures were diluted 100X in the media specified. Those conditions include standard media conditions as well as single component changes: 0% SDS, 0.01% SDS, 0.02% (wt/v) Arabinose, 0 mM Tris-HCl, 30 µg/mL Chloramphenicol. The cultures were grown in triplicate for each condition and in a SpectraMax i3 plate reader, shaking at 34 °C for 24 h. The $OD_{600}$ was measured about every 5 mins. The doubling times were then calculated as previously described.

*Bacterial and mammalian* co*-culturing*

HEK293T cells containing one copy of mCherry marker (red) integrated into the AAVS1 locus, were grown at 40-50% confluency in DMEM high glucose medium (Thermo Fisher cat# 11965175) with 10% inactivated Fetal Bovine Serum (FBS Thermo Fisher cat# 10082147), 100X MEM NEAA (non-essential amino acids Thermo Fisher cat# 11140050), and 100X diluted anti-anti cocktail (Antibiotic-Antimycotic - 10,000 units/mL of penicillin, 10,000 µg/mL of streptomycin, and 25 µg/mL of Gibco Amphotericin B - Thermo Fisher cat# 15240112). Commercially acquired *E. coli* DH5α bacteria were used as control to the *E. coli* DEP *mutS*$^{-}$ or DEP* strain. A plasmid containing Clover (green marker) containing a UAA stop codon compatible with the biocontained strain DEP, and under the selection marker ampicillin was transformed into both DH5α and DEP* strains in order to visualize them with the mammalian cells (red). BipA-dependent auxotroph DEP* bacteria were grown to OD 0.6, in LB medium supplemented with 1% L-arabinose, 100 µM BipA, 100 ug/ml carbenicillin and 25ug/ml chloramphenicol and then washed 3 times with 1X PBS. DEP* culture conditions with L-arabinose, carbenicillin, and chloramphenicol supplements did slightly affect HEK293T early cell growth compared to untreated cells, though insufficient to affect conclusions drawn from these experiments. DH5α strain was grown to OD 0.6 with 100ug/ml carbenicillin. The pellet of 10-milliliter bacterial cell culture was re-suspended in mammalian cell medium as described above without any antibiotics and anti-anti, and split equally among all conditions and their replicates. Auxotroph bacteria are added to HEK293T cells plated in pre-treated 12-well plates in 2 ml mammalian cell medium. The co-culture is incubated overnight before media with bacterial cells is removed and HEK293T cells are washed three times with 1X PBS (phosphate buffered saline Thermo Fisher cat# 10010023) and replenished with fresh media as conditions indicate. Media was replaced and added fresh to all conditions daily for 7 days. Imaging cells was done with the inverted microscope Nikon Eclipse TS100 at Day 2, Day 4, and Day 7 post initial co-culture at 200X magnification.

Conditions:

Control: HEK293T grown in regular 10% FBS media with anti-anti and NEAA as described above.

DH5α: HEK293T cells co-cultured with this strain in mammalian cell media supplemented with 100ug/ml carbenicillin to maintain plasmid during growth, and absence of anti-anti.

DH5α; anti-anti (antibiotic cocktail): HEK 293T cells co-cultured with this strain in mammalian cell media supplemented with 100ug/ml carbenicillin to maintain plasmid during growth, and presence of anti-anti cocktail.

DH5α; anti-anti after Day 2: HEK 293T cells co-cultured with this strain in mammalian cell media supplemented with 100ug/ml carbenicillin to maintain plasmid during growth, and absence of anti-anti cocktail. At 48 hours anti-anti added and maintained to Day 7.

DH5α; anti-anti; no anti-anti after Day 2: HEK 293T cells co-cultured with this strain in mammalian cell media supplemented with 100ug/ml carbenicillin to maintain plasmid during growth, and presence of anti-anti until Day 2. After Day 2 no anti-anti added and maintained to Day 7.

DEP*: HEK 293T cells co-cultured with the biocontained strain in media supplemented with L-arabinose, 25ug/ml chloramphenicol and 100ug/ml carbenicillin to maintain bacteria and green marker. No bipA or anti-anti added.

DEP*; bipA: HEK 293T cells co-cultured with the biocontained strain in media supplemented with L-arabinose, 25ug/ml chloramphenicol and 100ug/ml carbenicillin to maintain bacteria and green marker. 100 uM bipA and no anti-anti added.

DEP*; bipA after Day 2: HEK 293T cells co-cultured with the biocontained strain in media supplemented with L-arabinose, 25ug/ml chloramphenicol and 100ug/ml carbenicillin to maintain bacteria and green marker. No bipA or anti-anti added. At 48 hours bipA at 100uM concentration added and maintained to Day 7.

DEP*; anti-anti: HEK 293T cells co-cultured with the biocontained strain in media supplemented with anti-anti, L-arabinose, 25ug/ml chloramphenicol and 100ug/ml carbenicillin to maintain bacteria and green marker. No bipA added.

DEP*; bipA; anti-anti: HEK 293T cells co-cultured with the biocontained strain in media supplemented with anti-anti, L-arabinose, 25ug/ml chloramphenicol and 100ug/ml carbenicillin to maintain bacteria and green marker. 100uM bipA added.

*Persistence*

Persistence was evaluated by two kinds of assays: Plate reader and colony count. For the plate reader case, DEP, DEP.e3, DEP.e4, and DEP.e5 cultures were grown overnight in permissible media conditions with 100 μM BipA. For cells harvested at mid-exponential phase, the cultures were diluted 100X and grown to that state. Both stationary phase and mid-exponential phase cultures were then washed twice with LB media and resuspended in the original volume of non-permissible media containing all specified media components except BipA. The resuspended cultures were then diluted 100X into non-permissible media in triplicate for each time point to be tested. The specified concentration of BipA was then added back to those cultures at the specified time points. Typically, the BipA re-addition occurred at 10 μM or 5 μM concentrations and at hourly or daily intervals. The cultures were then incubated with shaking in SpectraMax i3 plate readers in a flat, clear bottom 96-well plate with breathable and optically transparent seal for an upwards of 84 hours at 34 °C. Approximately every five minutes the $OD_{600}$ was measured to determine cell growth kinetics.

Colony count assays used resuspended cultures that were generated using the same protocol as that for the plate reader persistence assay. These cultures were then grown in non-permissible media at 34 °C and shaking at 250 r.p.m in 10 mL culture tubes. At the specified time point, the cultures were removed and 10 μL of culture was serially diluted to a $10^8$-fold dilution. The dilutions were then plated on permissible solid media containing 10 μM or 5 μM BipA. The plates were then incubated at 37 °C for 48 h. The cultures were grown in triplicate unless otherwise specified.

### SI Figures

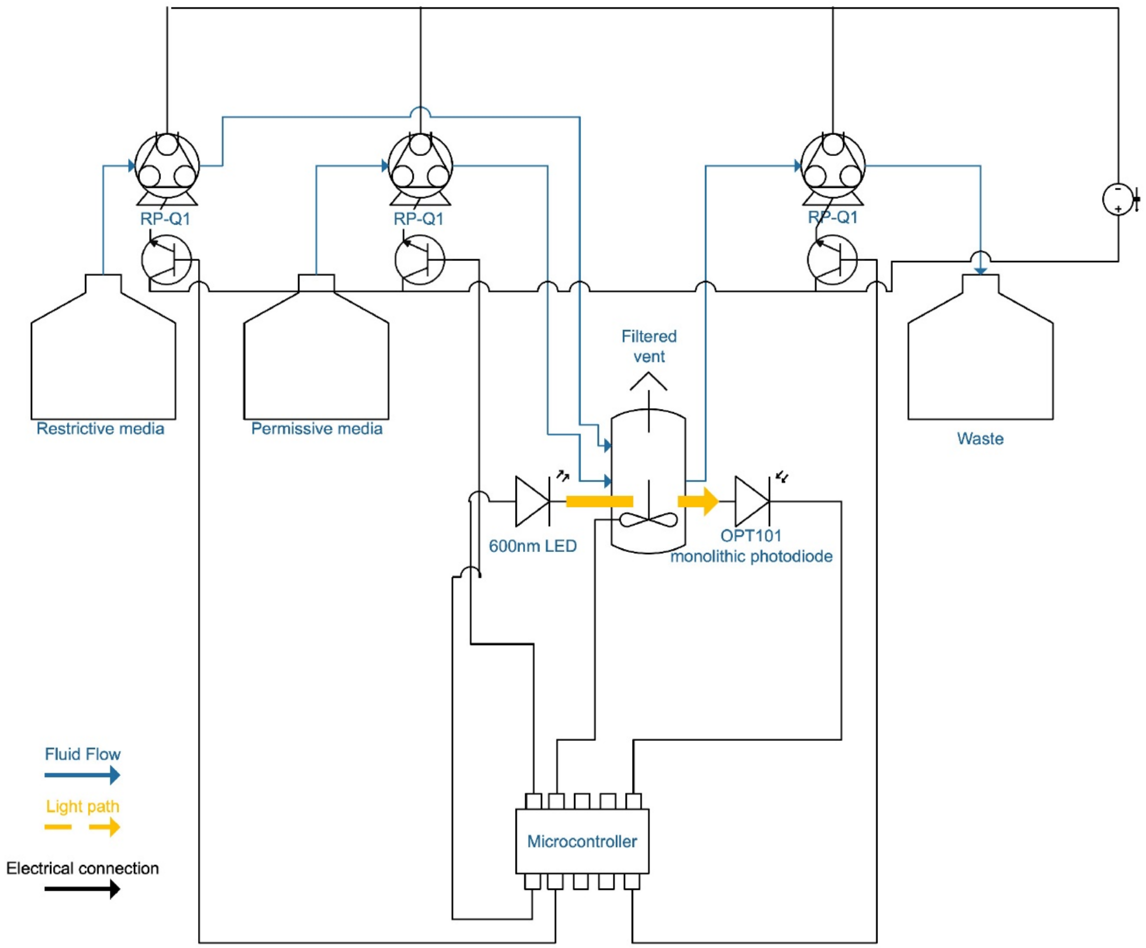


**Figure S1.** Fluid and electrical flow diagram of custom chemostats used for automated continuous evolution.

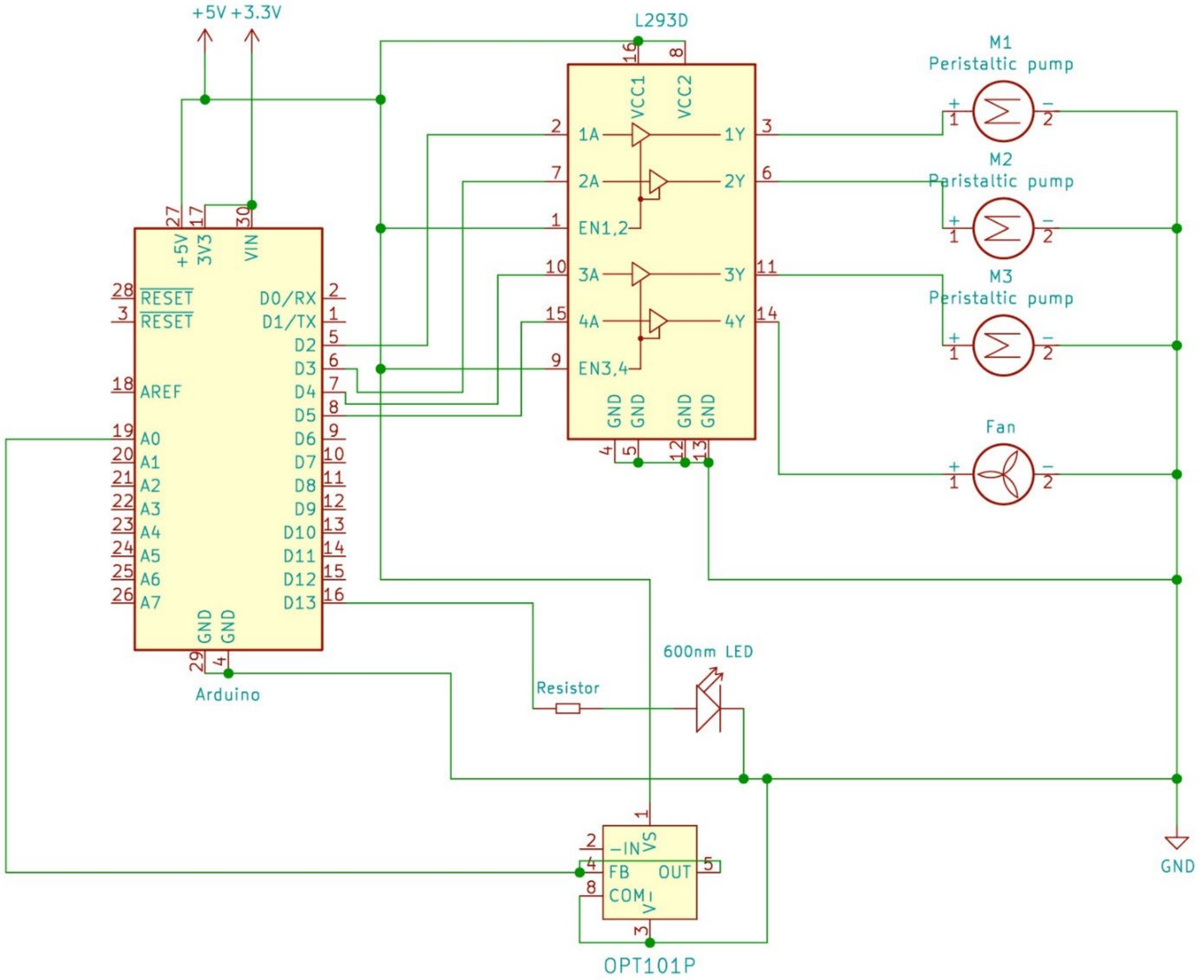


**Figure S2.** Chemostat circuit schematic.

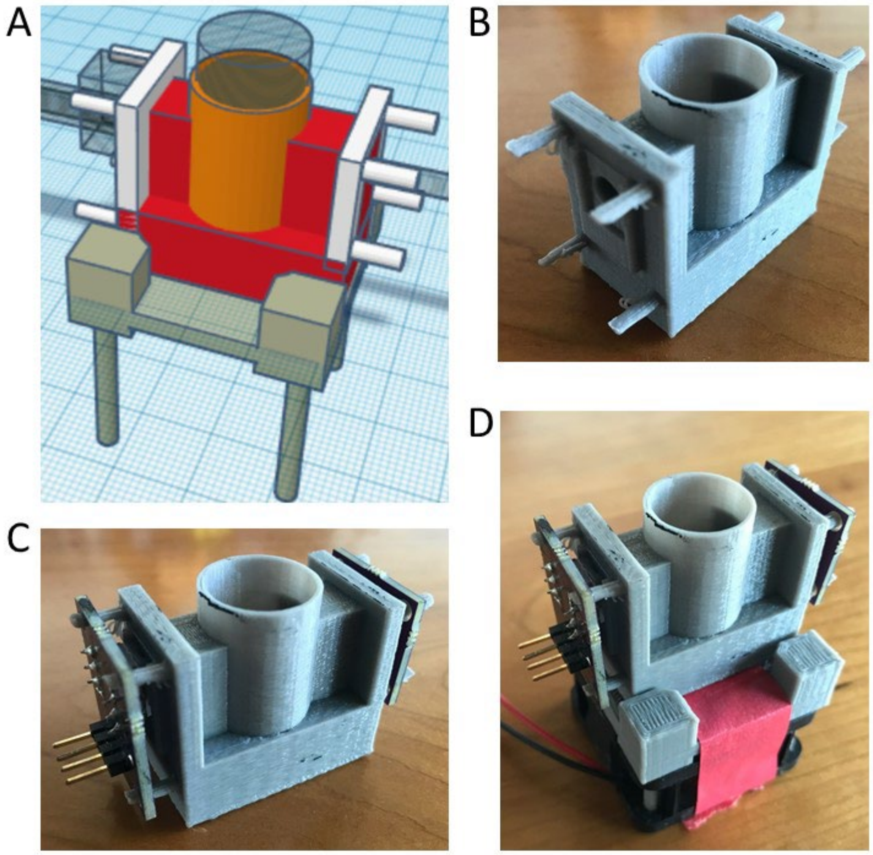


**Figure S3.** Representative 3D-printed chemostat sleeve stand. (A) CAD-rendering. (B) Actual stand, alone. (C) Stand with mounted LED and photodiode components. (D) Assembled sleeve with stir fan mount.

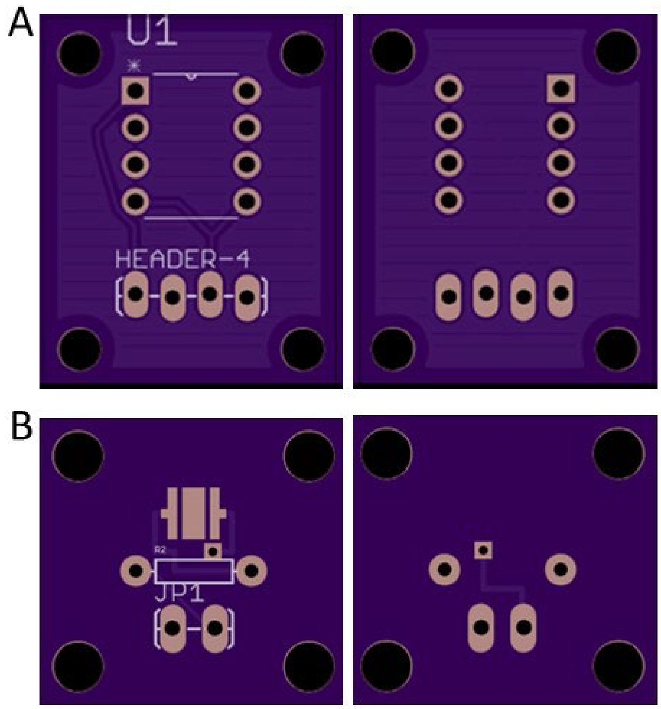


**Figure S4.** Custom PCBs used to create light path.

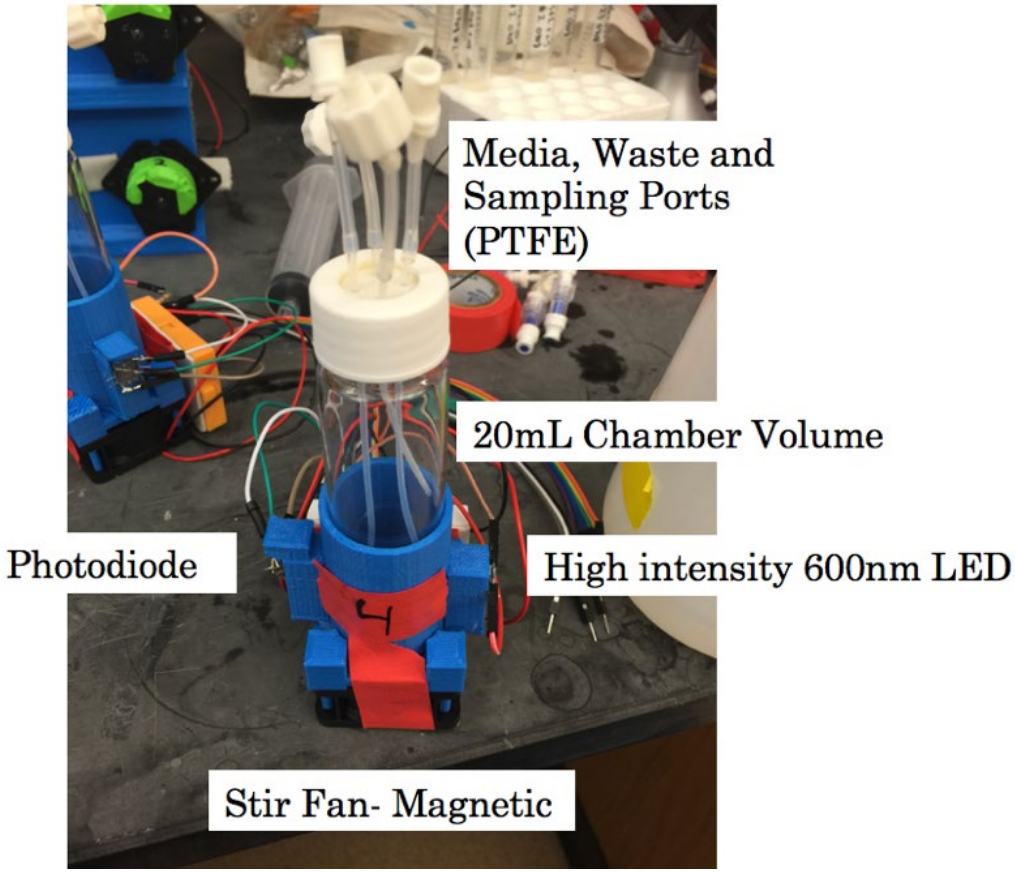


**Figure S5**. Representative chemostat smart sleeve used for automated continuous evolution of synthetic auxotrophs. Important features are labeled, and pumps are in the background.

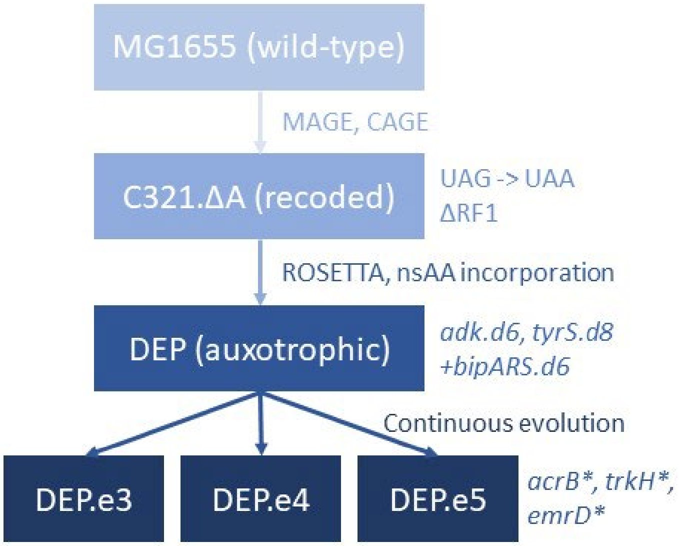


**Figure S6.** Ancestry of synthetic auxotrophs discussed in this manuscript.

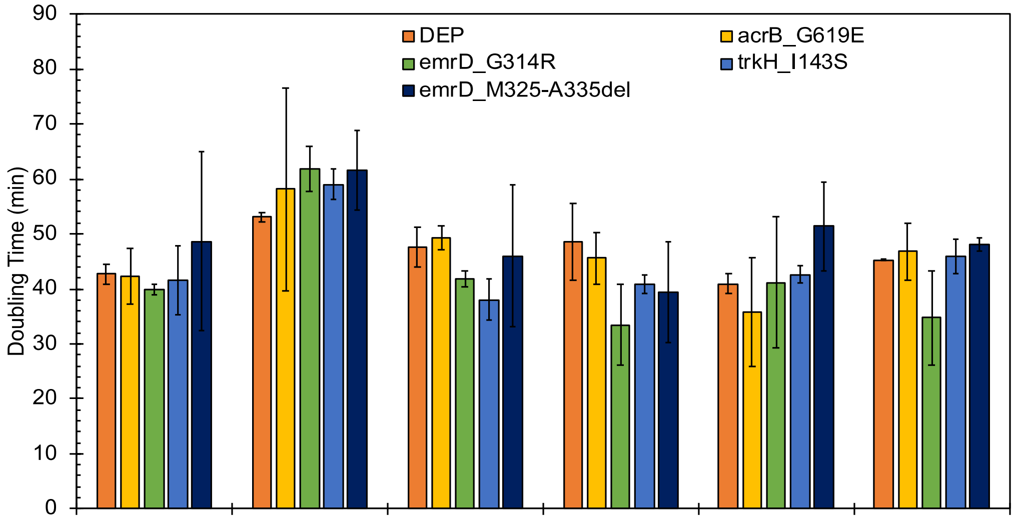


| | | | | | | |
|---|---|---|---|---|---|---|
| 0% SDS | - | + | - | - | - | - |
| 0.005% SDS | + | - | - | + | + | + |
| 0.01% SDS | - | - | + | - | - | - |
| 0.02% (wt/v) Arabinose | - | - | - | + | - | - |
| 0.2% (wt/v) Arabinose | + | + | + | - | + | + |
| 0 mM Tris | - | - | - | - | + | - |
| 20 mM Tris | + | + | + | + | - | + |
| 15 ug/mL Chloramphenicol | + | + | + | + | + | - |
| 30 ug/mL Chloramphenicol | - | - | - | - | - | + |

**Figure S7.** Effect of varying media components whose transport may be altered by alleles discovered from evolution on cellular doubling times, with no substantial differences observed.

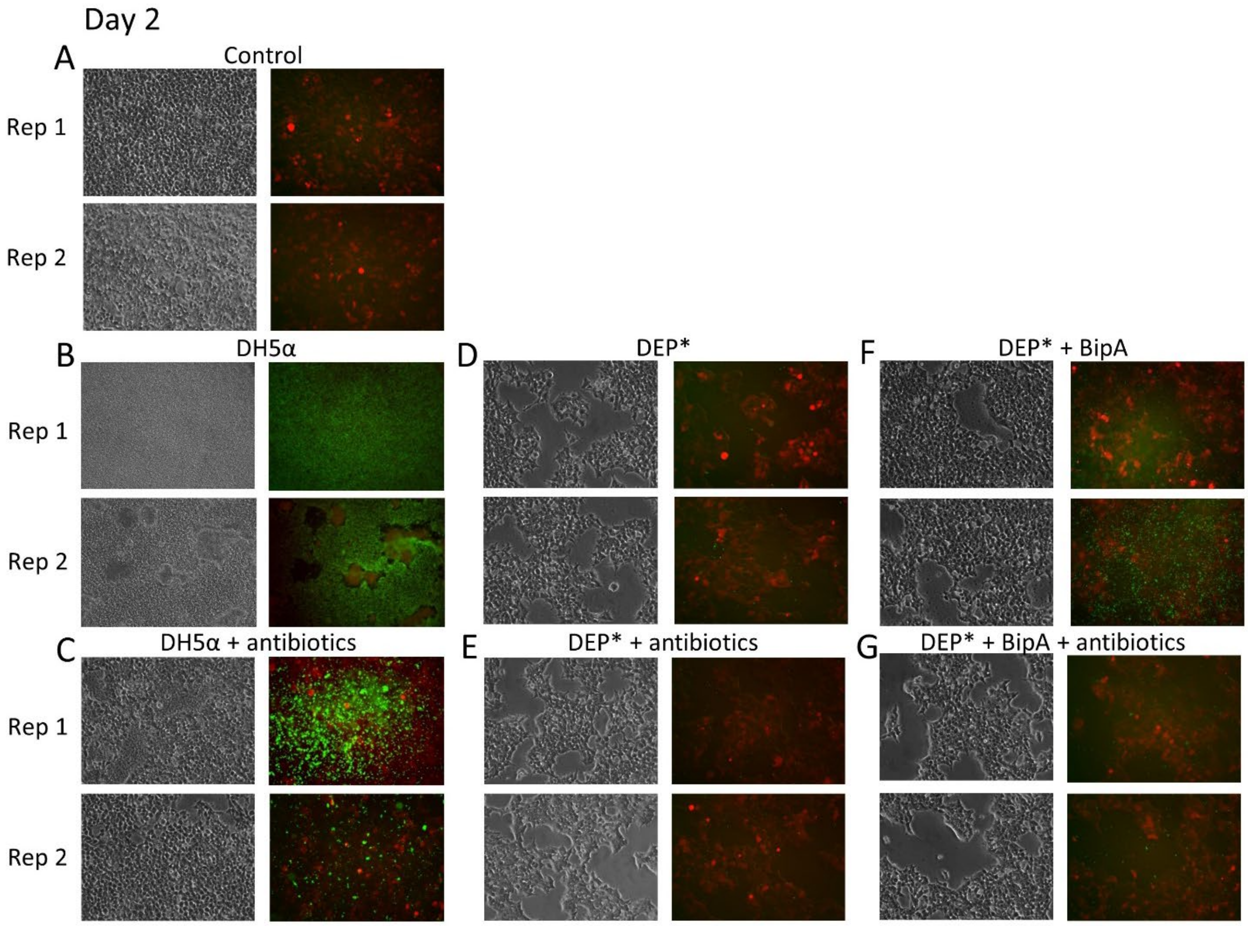


**Figure S8.** Imaging replicates of Day 2 conditions in Figure 4 of the main text. Note that the two replicates shown are in addition to the third replicate that is included in Figure 4. Additional conditions include Figure S8E where DEP* is co-incubated with HEK293T in the absence of BipA and in the presence of antibiotic cocktail, and Figure S8G where DEP* is co-incubated with HEK293T cells in presence of BipA and antibiotic cocktail.

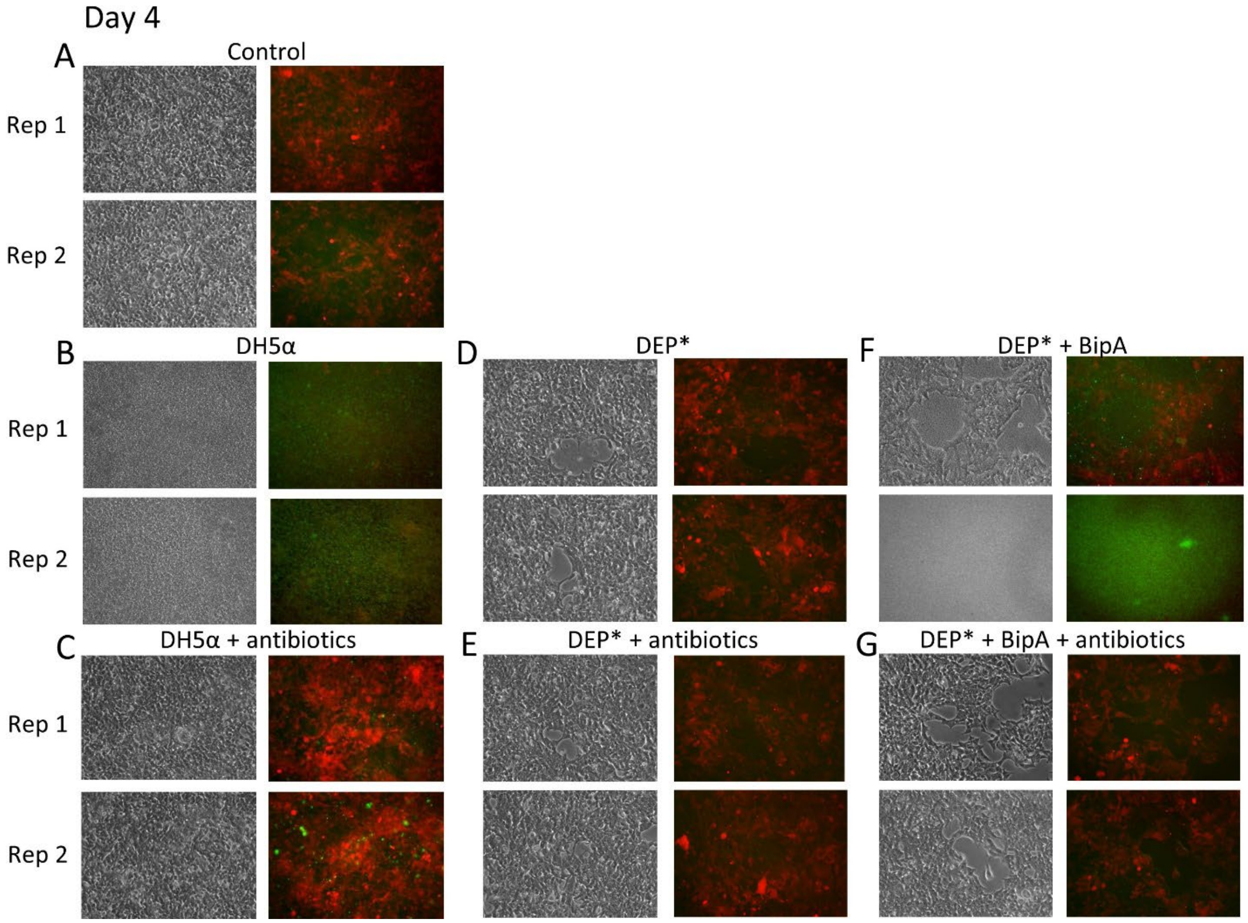


**Figure S9.** Imaging replicates of Day 4 conditions in Figure 4 of the main text, with additional conditions as described in Figure S8.

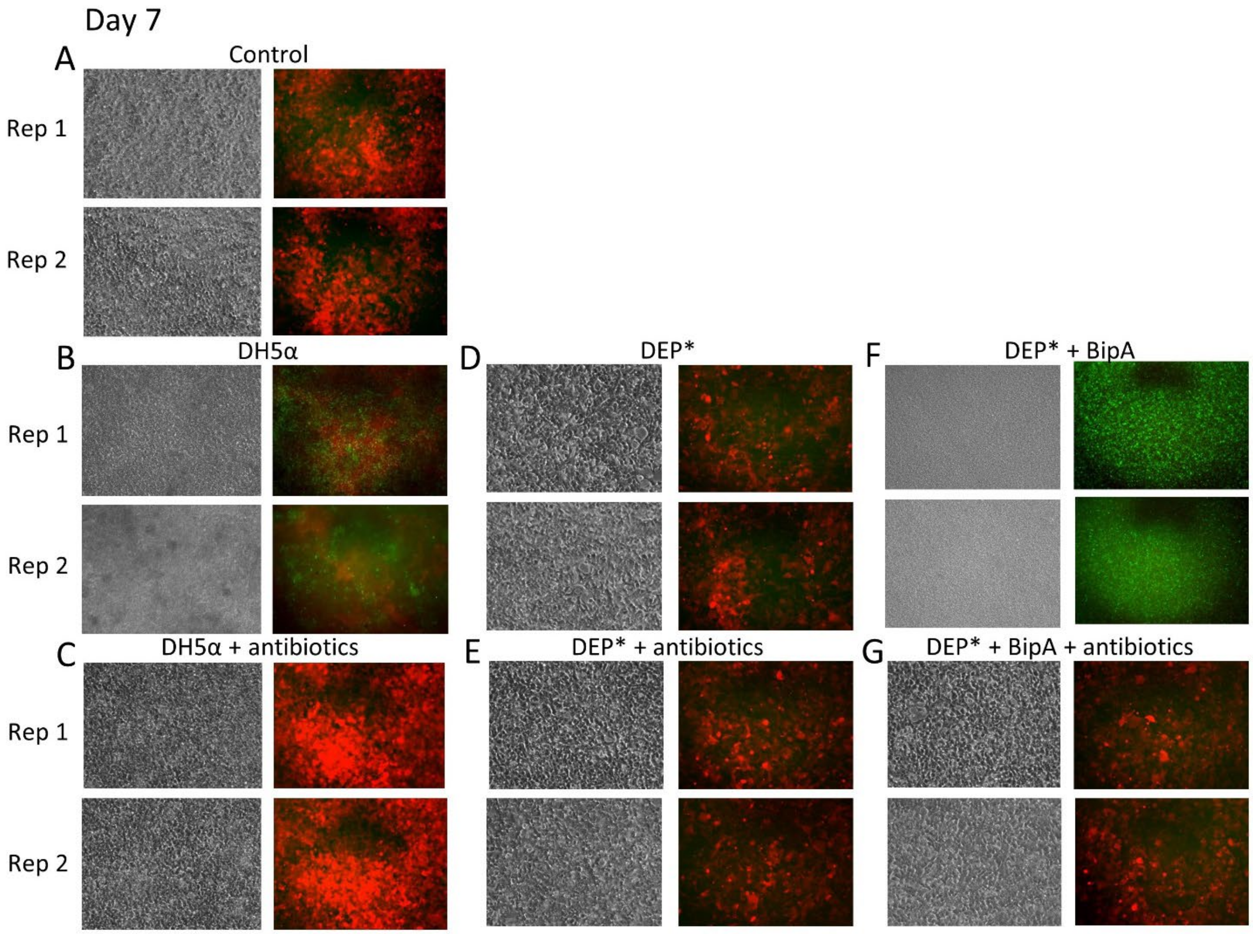


**Figure S10.** Imaging replicates of Day 7 conditions in Figure 4 of the main text, with additional conditions described in Figure S8.

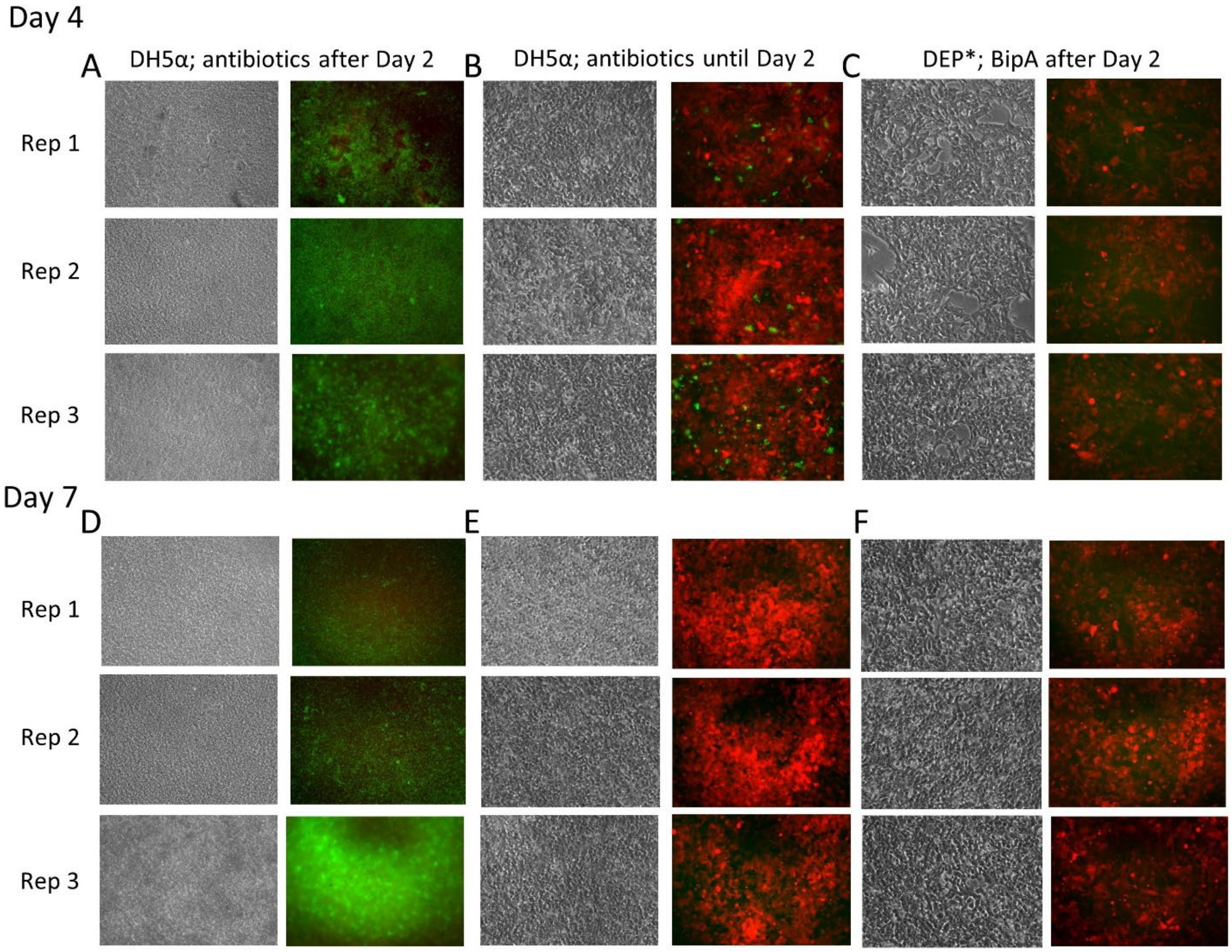


**Figure S11.** Investigation of persistence of bacteria in mammalian co-culture based on morphology. (A) Day 4. HEK 293T cells co-cultured with DH5α in absence of anti-anti for 2 days, then addition of anti-anti until Day 7. (B) Day 4, HEK 293T and DH5α cultured in presence of anti-anti for 2 days before removing anti-anti for the duration of experiment. (C) Day 4. DEP* maintained in absence of BipA media for 2 days, then BipA added at Day 2-7 to investigate possible persistence and recovery of bacterial cells. (D) Day 7 of (A). (E) Day 7 of B. (F). Day 7 of (C). Note that Figure 4 in the main text displays one replicate from each of panels C and F.

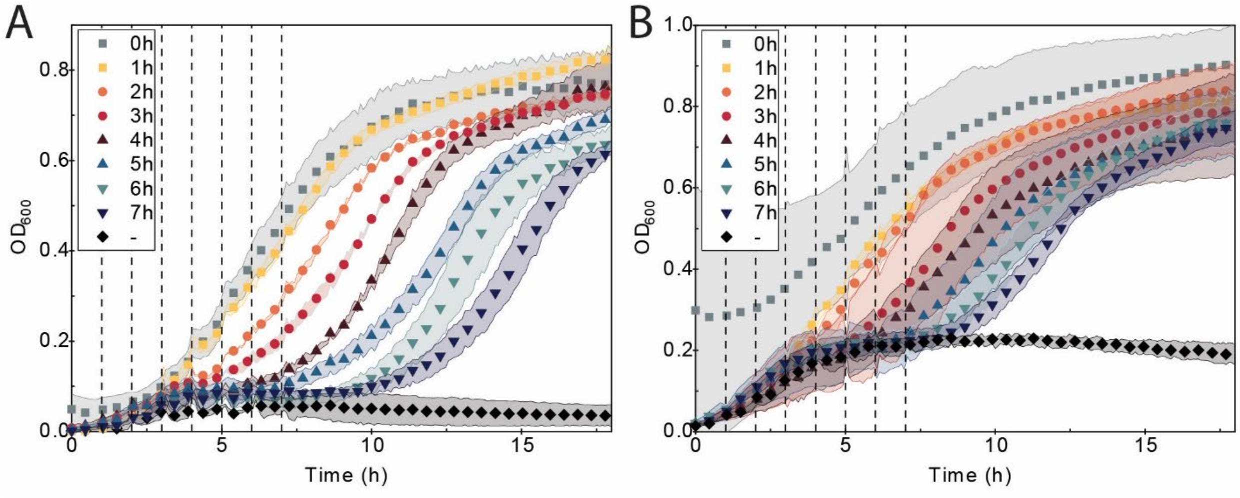


**Figure S12.** Rescue of DEP following deprivation and subsequent re-addition of BipA at the specified time point. The ability to recover growth provides another mechanism to control the biocontained strain. At the specified time point, 10 μM BipA was added to the specified well. (A) Rescue with an initial culture at mid-exponential phase. (B) Rescue with an initial culture at stationary phase.

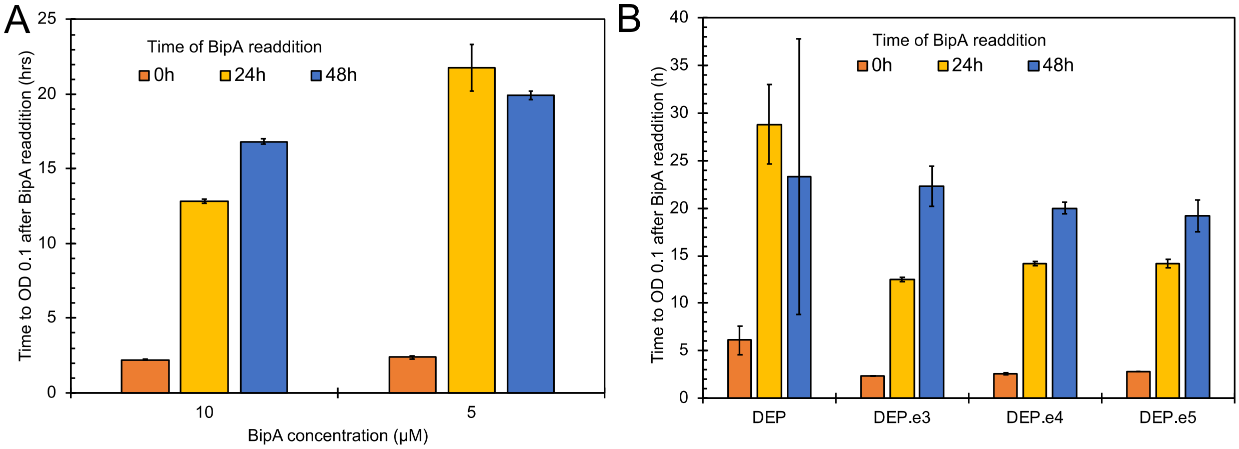


**Figure S13.** Demonstration of rescue for DEP and the evolved strains up to 48hrs after BipA removal. Despite no substantial growth following BipA removal, the cultures were able to be rescued following BipA addition after 24 and 48 h. Growth is measured and indicated by the time it takes the cultures after BipA re-addition to reach an $OD_{600}$ of 0.1 and maintain it for at least 30 minutes. (A) Growth following re-addition of different concentrations of BipA to DEP cultures. (B) Growth following re-addition of 5 μM BipA to DEP, DEP.e3, DEP.e4, and DEP.e5 cultures.

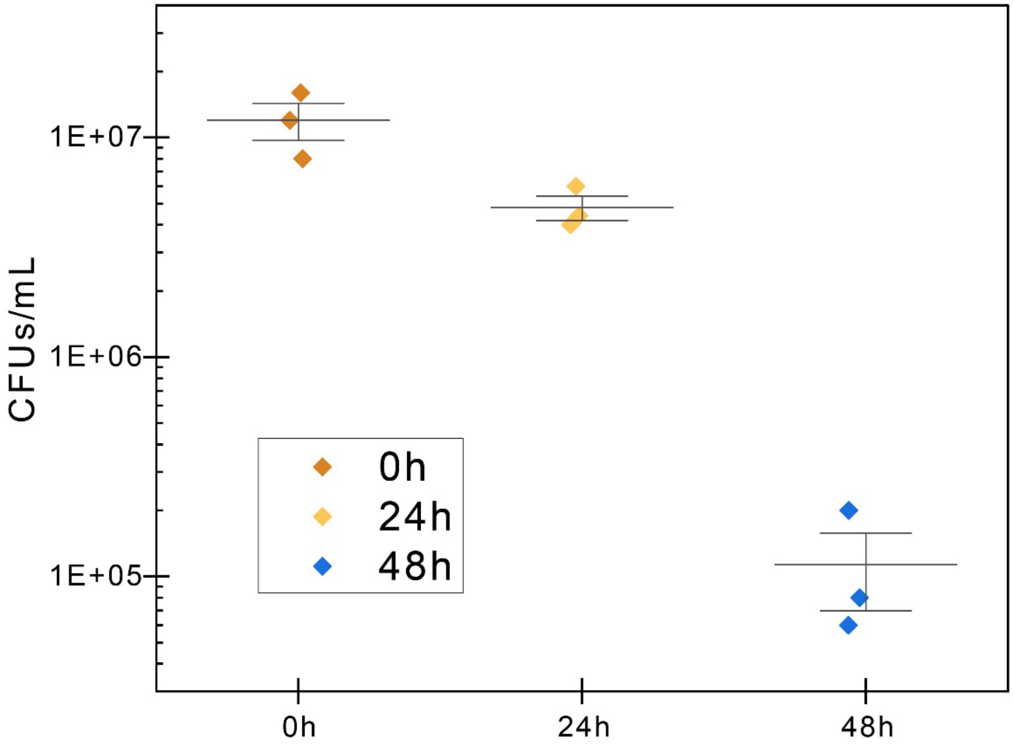


**Figure S14.** Direct measurement of bacterial persistence by monitoring density of viable colony forming units (CFUs/mL) present in cultures at times indicated after BipA removal. Horizontal line indicates average and vertical error bars indicate 1 standard deviation based on the three replicates shown. Samples were also collected at 72 h and 96 h. Volumes as high as 5 µL of the original culture were plated on permissive media (assay lower detection limit of 200 CFUs/mL) and allowed to incubate for up to 48 hours. No CFUs were observed from the 72 h and 96 h time points.

### SI Tables

**SI Table 1.** Transporter mutations observed from evolution of DEP. Steric clash predicted upon substitution is indicated in red.

| | Description | Visualization |
|---|---|---|
| AcrB_G619E | AcrB has been implicated in the transport of several different classes of antimicrobial molecules, including SDS and chloramphenicol, through its formation of a homotrimer in a three-part protein complex with AcrA and TolC[11–13]. While AcrB is non-essential[14], the deletion of *acrB* substantially affects sensitivity to antimicrobial agents. The $\text{AcrB}^{G619E}$ variant observed occurs in a switch-loop region that separates the access pocket from the deep pocket within the periplasmic portion of the protein. The switch-loop may participate in high-traffic export mechanisms[15,16]. AcrB is known to transport planar aromatic cations that resemble BipA via a channel in which the switch-loop plays a role[17]. Despite this fact, $\text{AcrB}^{G619P}$, a mutation of the same residue we observed, was found to affect drug resistance only minimally in a mutagenesis study that targeted the switch-loop[18]. | 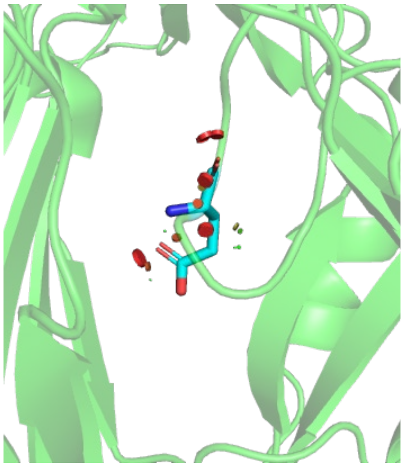 PDB ID: 4DX5 [16] |
| EmrD_G314R | The two EmrD mutations observed affect the $10^{th}$ and $11^{th}$ helices of this protein, which have been implicated as a major sub-pathway for substrate transport[19]. These mutations may cause disruption to efflux function as molecular dynamics simulations have suggested critical hydrogen bonding across these transmembrane helices to stabilize the structure during transport. Alterations to these transporters might also alter levels of arabinose in the cytosol[20]. While the biphenylalanine aminoacyl-tRNA synthetase in DEP is arabinose-regulated, an increase in intracellular concentration is unlikely to increase expression given that induction is binary and often saturated at these concentrations[21,22]. | 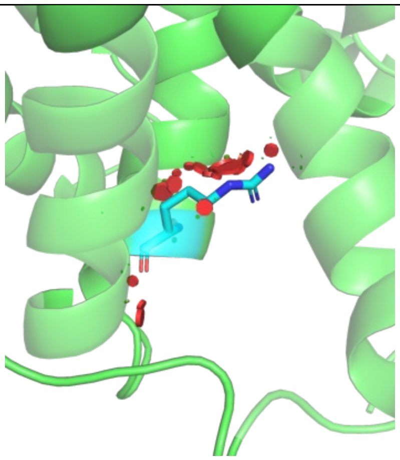 PDB ID: 2GFP [23] |

| | | |
|---|---|---|
| TrkH_I143S | TrkH is a potassium ion transporter that contains 10 transmembrane helices that allow for transport from the periplasm to the cytoplasm[24]. Based on a homology model, the TrkH variant discovered here may perturb the helical structure of one of the transmembrane helices thus disrupting the structure and function of the protein. | 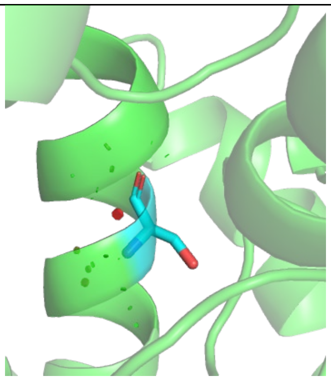 Structure based on a homology model produced based on PDB ID 6V4L [25] |

**SI Table 2.** Complete list of variants obtained from DEP evolution.

| Emergent Mutation | Variant e3 | Variant e4 | Variant e5 |
|---|---|---|---|
| 481597 (acrB) | ✓ | ✓ | ✓ |
| 813092 (150 bp upstream of N gene) | ✓ | | |
| 890773 (6 bp downstream of mdfA) | ✓ | | |
| 1438014 (pinR) | ✓ | | |
| 1438003 (pinR) | | ✓ | |
| 1438009 (pinR) | | | ✓ |
| 2163524 (mdtC) | ✓ | | |
| 3856781 (emrD) | ✓ | | |
| 4035491 (trkH) | ✓ | ✓ | ✓ |
| 3993912 (cyaA) | | | ✓ |
| 3856814 (emrD) | | ✓ | ✓ |
| 4041964 (rrlA) | ✓ | | ✓ |
| 4040603 (rrlA) | | ✓ | |
| 4040604 (rrlA) | | | ✓ |
| Reversion of Mutation from DEP | | | |
| 707016 (210 bp upstream glnS; 365 bp downstream chiP) | | × | × |
| 707062 (260 bp upstream glnS; 320 bp downstream chiP) | × | × | |
| 816377 (49 bp upstream tetR; 36 bp downstream bla) | × | | |
| 817324 (52 bp upstream bla; 165 bp downstream bioF) | | | × |
| 1299471 (72 bp downstream tdk) | | | × |
| 1300208 (49 bp upstream tdk) | | × | |
| 1438012 (pinR) | × | × | × |
| 2861919 (mutS) | | × | × |
| 4040606 (rrlA) | × | × | × |

**SI Table 3.** Build of materials for custom chemostat.

| BOM | Part Description | # | Part # | Notes | Supplier |
|---|---|---|---|---|---|
| Pump block | RP-Q1 | 3 | RP-Q1.2N-P20A-DC3 | For small volumes (<10mL working volume) | Takasago Fluidics |
| | or | | | | |
| | RP-CIII1 | 3 | RP-CIII1.6S-2Z-DC5V | For larger volumes (50mL vials) | Takasago Fluidics |
| | L239D | 1 | L293D | Quad half-h motor drivers | Digikey or similar |
| | Stand | 1 | n/a | 3D printed | |
| Power | 5V 2A DC power supply | 1 | generic | | Adafruit or similar |
| Media reservoirs | 2L sterilizable bottles | 3 | generic | 2 for media, one for waste. Can use any size | |
| | "octopus" modified lids | 3 | generic | see sketch for adaptation. Seal openings around tubing with autoclavable silicone adhesive | |
| Vials | 15mL HPLC glass vial | 1 | | | Millipore Sigma or similar |
| | 15mL vial lid with PTFE/silicone septum (PP or heat tolerant) | 1 | | | Millipore Sigma or similar |
| | rigid 1mm ID PTFE tubing | 4 | | insert through septum, 2 lines terminating near top of the vial (media), 2 lines at around halfway point (waste/sampling lines) | McMaster Carr |
| | 7mm x 2mm stir bar | 1 | 14-513-63 | generic is fine as long as it fits | Fisher Scientific |
| | JB weld twin tube | 1 | | Cover pierced septum with JB weld, using foil to contain it while it sets | Amazon |
| Fan block | 40mm x 40mm 5v computer fan | 1 | | | Amazon |
| | 5mm disk magnet | 2 | | glue magnets to fan in +/- orientation | Amazon |
| | epoxy | 1 | | | Amazon |
| | fan adapter | 1 | | glue vial sleeve to fan adapter, slot fan into guides | |
| Vial sleeve | 600nm LED | 1 | Recommend L1C1-AMB1000000000 or XPEAMB-L1-0000-00301 | Doesn't need PCB, but makes soldering/fitting easier | Newark |
| | Photodiode | 1 | OPT101P | Doesn't need PCB, but makes soldering/fitting easier | Newark |
| | vial adapter | 1 | | 3d printed, use guides to slot in photodiode and LED on each side | |
| | Through hole resistor, 3.3K | 1 | | Adapt resister value for LED intensity | |
| | PCB for LED/photodiodes | 2 | | OSH park | |
| Tubing | 1/32" ID (1mm OK) silicone tubing | 50 ft | | Cut as needed to connect per diagram | Cole Palmer |
| | 3/32" Male Luer to Barb | 50 | | As needed for assembly | Cole Palmer |
| | 3/32" Female Luer to Barb | 50 | | As needed for assembly | Cole Palmer |

| | | | | | |
|---|---|---|---|---|---|
| | Luer Male plug | 50 | | As needed for assembly / capping of sample lines | Cole Palmer |
| | Sterile Luer compatible .22uM syringe filters, 25mm PVDF | 4 | EW-06060-63 | One for each reservoir/waste for gas vent, one for vial for gas exchange | Cole Palmer |
| | 3/16" ID silicone tubing | 25 ft | | Protects the smaller tubing as it passes through the lids to the media reservoirs | Cole Palmer |
| | 1/4" Male Luer | 25 | | | Cole Palmer |
| Micro-controller | Arduino or Arduino compatible microcontroller | 1 | 5V pro micro | Any microcontroller with a 5V tolerating ADC pins will do for ease of use, if careful with voltages a more modern 3.3V microcontroller will do, and will allow for more interesting / easy designs like a circuit python datalogger that can stand alone without a computer | Adafruit |
| | RPi or laptop | 1 | | Controls uC logic | Adafruit |

**SI Table 4.** Oligonucleotides used in this study.

| Oligo Name | Sequence |
|---|---|
| acrB-MAGE | C*A*ACGTTGAGTCGGTGTTCGCCGTTAACGGCTTCGGCTTTGCGGAACGTGGTCAGAATACCGGTATTGCGTTCGTTTCCTTGAAGGACTG |
| trkH-v2-MAGE | C*C*CACCCACGCCGAGGATAGGCAGTATCGCAACCGCCAACACGCTGATCCCCATCCCGCCAAACCATTGCAGCATCTGGCGATAAAAGAG |
| emrD-MAGE | G*T*CTGGACGCTGCTCGTTCCCGCCGCGCTGTTCTTTTTCGGTGCCAGGATGCTGTTTCCGCTGGCGACCAGCGGCGCGATGGAGCCGTTC |
| acrB-mut-Fv2 | CTTCGGCTTTGCGGA |
| acrB-wt-Fv2 | CTTCGGCTTTGCGGG |
| acrB-Rv2 | GCATCTTTGATTTGCGAGAAAG |
| trkH-v3-mut-F | CGGGATGGGGATCAG |
| trkH-v3-wt-F | CGGGATGGGGATCAT |
| trkH-v3-R | CGAAATAACCGATACTGGC |
| emrD-mut-F | GCTGTTCTTTTTCGGTGCCA |
| emrD-wt-F | GCTGTTCTTTTTCGGTGCCG |
| emrD-R | CTTCGGTGACAGCAGAGCTG |
| acrB-seq-F | GCATGGCCTATCTGTTCG |
| trkH-seq-F | GCTCCCTTTTATCTTCTCGG |
| emrD-seq-F | CAGTATTTTGTTTATTCTGCCG |
| acrB-seq-R | CTGAGTCAGTTTTTCGTGAC |
| trkH-seq-R | ATCGAAATAACCGATACTGGC |
| emrD-seq-R | GTGGGGGAAATTTTTAATTGCC |
